# Laboratory evolution reveals a two-dimensional rate-yield tradeoff in microbial metabolism

**DOI:** 10.1101/414912

**Authors:** Chuankai Cheng, Edward J. O’Brien, Douglas McCloskey, Jose Utrilla, Connor Olson, Ryan A. LaCroix, Troy E. Sandberg, Adam M. Feist, Bernhard O. Palsson, Zachary A. King

## Abstract

Growth rate and yield are fundamental features of microbial growth. However, we lack a mechanistic and quantitative understanding of the rate-yield relationship. Studies pairing computational predictions with experiments have shown the importance of maintenance energy and proteome allocation in explaining rate-yield tradeoffs and overflow metabolism. Recently, adaptive evolution experiments of *Escherichia coli* reveal a phenotypic diversity beyond what has been explained using simple models of growth rate versus yield. Here, we identify a two-dimensional rate-yield trade-off in adapted *E. coli* strains where the dimensions are (A) a tradeoff between growth rate and yield and (B) a tradeoff between substrate (glucose) uptake rate and growth yield. We employ a multi-scale modeling approach, combining a previously reported coarse-grained small-scale proteome allocation model with a fine-grained genome-scale model of metabolism and gene expression (ME-model), to develop a quantitative description of the full rate-yield relationship for *E. coli* K-12 MG1655. The multi-scale analysis resolves the complexity of ME-model which hindered its practical use in proteome complexity analysis, and provides a mechanistic explanation of the two-dimensional tradeoff. Further, the analysis identifies modifications to the P/O ratio and the flux allocation between glycolysis and pentose phosphate pathway as potential mechanisms that enable the tradeoff between glucose uptake rate and growth yield. Thus, the rate-yield tradeoffs that govern microbial adaptation to new environments are more complex than previously reported, and they can be understood in mechanistic detail using a multi-scale modeling approach.

## 1 INTRODUCTION

Growth rate and yield are basic features of microbial life that are widely implicated in cell fitness, adaptation, and evolution (Lipson, 2015). The specific growth rate *µ* represents number of doublings of bacterial density per unit time (Monod, 1949). The yield, *Y*, is the ratio between the biomass dry weight produced and the weight of the substrate uptaken (Monod, 1949; Pirt, 1965). There is great interest in developing quantitative descriptions of the relationship between *µ* and *Y*. The wide-ranging measurements of *µ* and *Y* (Fig. 1A) across microbial communities and environments raised interest into the exact nature of the *µ*–*Y* relationship (Lipson, 2015). At low *µ*, positive correlations between *µ* and *Y* have been observed (Nanchen et al., 2006), and these can be explained by non-growth-associated cell maintenance requirements that make slow growth inefficient (Pirt, 1965). At high *µ*, negative correlations between *µ* and *Y* are observed (Basan et al., 2015), and in *Escherichia coli*, these can be explained by a tradeoff between metabolic efficiency and enzymatic efficiency that lead to decreased *Y* when at high *µ* (Novak et al., 2006; Pfeiffer et al., 2001). In particular, *E. coli* exhibits a tradeoff between respiration, which has higher energy yield per carbon substrate (more metabolically-efficient), and acetate fermentation, which requires less enzyme per carbon substrate (more proteome-efficient) (Basan et al., 2015). Lipson proposes a synthesis of these observations where positive *µ*–*Y* correlation at low *µ* and negative *µ*–*Y* correlation at high *µ* are different parts of a bell-shaped *µ*–*Y* curve (Lipson, 2015). However, recent experiments suggest that adaptation to new environments can modify the bell-shaped *µ*–*Y* tradeoff (LaCroix et al., 2015; Sandberg et al., 2014; Bachmann et al., 2013).

**FIGURE 1.**
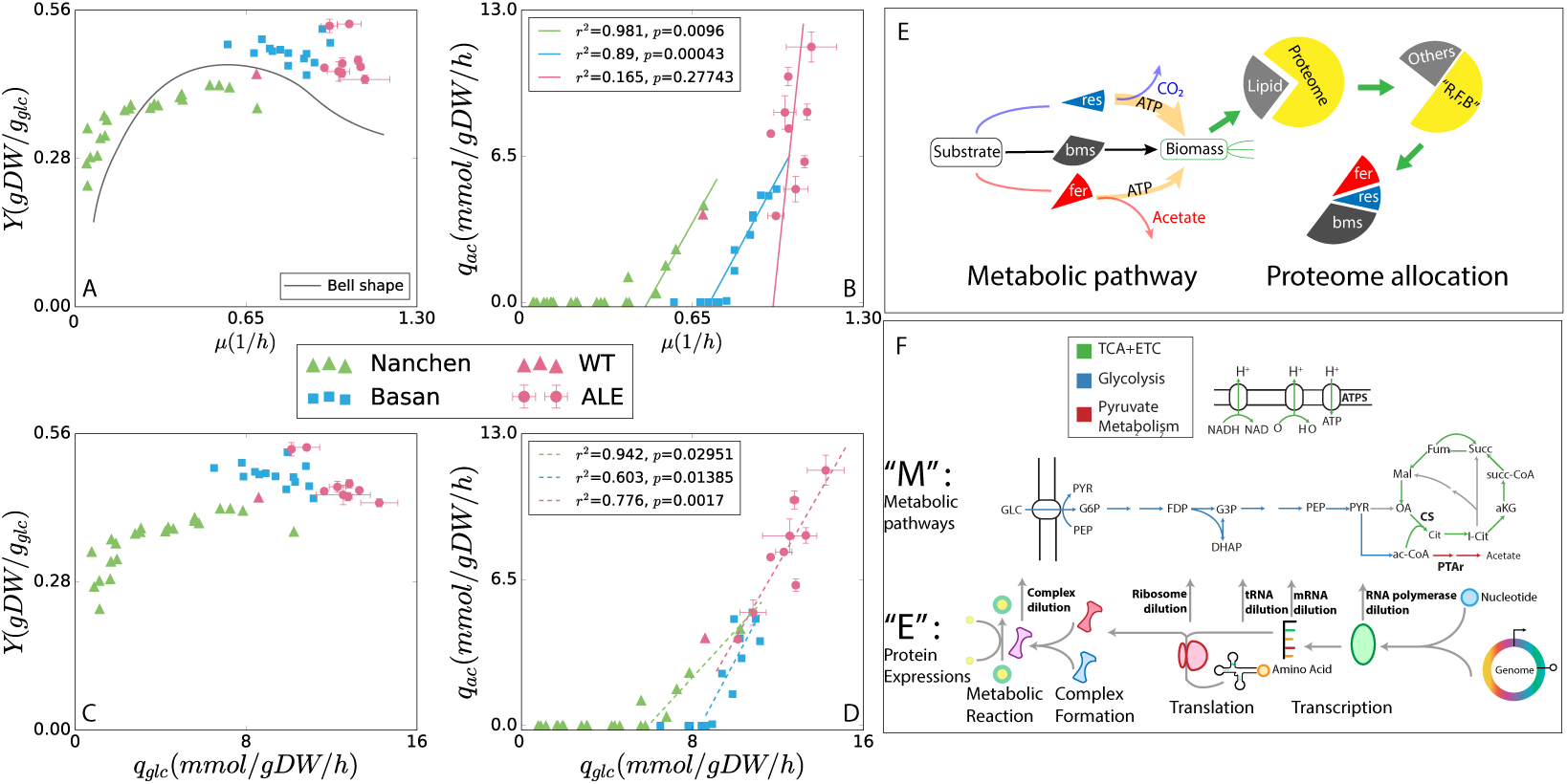
*E. coli* growth phenotypes (Data on the plots are recorded in Table S7–10 in Data (Expanded view).) in minimal media and multi-scale modeling approaches. (A–D) Phenotypic data for *E. coli* strains including *Y, µ, q*_*ac*_, and *q_glc_* data. Two datasets are presented from literature, for chemostat growth Nanchen et al. (2006) (green triangles) and substrate titration Basan et al. (2015) (blue squares). These are compared to strains adapted for maximum growth rate through ALE (this study; red circles; error bars for standard deviation across duplicates). The bell-shaped *µ*–*Y* relationship proposed by Lipson (2015) is included for reference. (E) Diagram of the SSME-model derived from Basan et al. (2015). The model consists of three pathways: respiration (res) and fermentation (fer) generate different amounts of energy, feeding the biomass (bms) pathway to synthesize biomass. (F) Diagram of the genome-scale ME-model that includes a genome-scale reconstruction of metabolic pathways and detailed protein expression machinery (O’Brien et al., 2013; Lloyd et al., 2018).

Microorganisms rapidly adapt to environmental niches (Booth, 2002; Elena and Lenski, 2003), and adaptation mechanisms can be studied directly through adaptive laboratory evolution (ALE) (Barrick and Lenski, 2013). When strains are adapted through ALE for growth in a liquid minimal medium, they achieve higher *µ* compared to the wild-type (Fig. 1A), ALE-adapted strains have been shown to rapidly acquire regulatory mutations that modify proteome allocation, but they do not acquire new metabolic capabilities within the time frame of reported short-term (4 to 8 weeks) adaptation experiments (LaCroix et al., 2015; Sandberg et al., 2014; Utrilla et al., 2016). By analyzing ALE-adapted strains, we can reveal the strategies that allow cells to optimize their proteome allocation for growth in an environmental niche, subject to the constraints of their metabolic capabilities (i.e. their repertoire of pathways) and constraints on the kinetic efficiencies of their enzymes (LaCroix et al., 2015; Utrilla et al., 2016; Ibarra et al., 2002).

Contrary to the expected negative *µ*–*Y* relationship at high *µ*, ALE experiments of *E. coli* selected for high *µ* in a minimal medium reveal an uncorrelated relationship between *µ* and *Y* (LaCroix et al., 2015; Sandberg et al., 2014). These experiments compared the phenotypes of highly-adapted, isogenic *E. coli* strains and revealed little variation in *µ* between strains but high variation in *Y*. Previous studies similarly reported that overflow metabolism can be nearly eliminated through genetic engineering without any effect on growth rate in *E. coli* (Peebo et al., 2014; Bekker et al., 2009). Thus, the negative *µ*–*Y* correlation at high growth rates does not appear to be a fundamental constraint on fast-growing cells. A mechanistic model of the full *µ*–*Y* relationship must be able to reconcile the bell-shaped curve observed for individual strains with the uncorrelated *µ*–*Y* phenotypes seen in ALE-adapted strains (Fig. 1A).

A number of theoretical and computational models have been developed to describe rate-yield tradeoffs. For the positive *µ*–*Y* correlation, maintenance requirements can be quantitatively described using algebraic growth laws (Pirt, 1965; Nanchen et al., 2006). This relationship can also be predicted for complete metabolic networks using genome-scale models (GEMs) of metabolism that encode non-growth associated maintenance (NGAM) costs in an optimization problem that can predict *µ* and *Y* when substrate uptake rates are known (Varma et al., 1993). For the negative *µ*–*Y* correlation, quantitative models of overflow metabolism have been developed (Basan et al., 2015; Mori et al., 2016; Molenaar et al., 2009). In particular, quantitative measurements of *E. coli* growth in well-controlled environments revealed a linear-threshold response of acetate excretion (*q_ac_*) with increasing *µ* (Basan et al., 2015). To represent the full range of the *µ*-*Y* relationship, a constraint allocation flux balance analysis model (CAFBA) was reported that combines a GEM with proteome allocation constraints (Mori et al., 2016). A similar solution can be formulated from a bottom-up reconstruction of metabolism and macromolecular expression (ME-model,O’Brien et al. (2013)) that incorporates the protein synthesis pathways into a GEM and applies coupling constraints related to enzyme kinetics parameters on each individual reaction. However, none of these models have been used to explain experiments where *µ* and *Y* are decoupled through laboratory evolution or genetic engineering (LaCroix et al., 2015; Sandberg et al., 2014; Peebo et al., 2014).

In this study, we show that the wide range of *µ*–*Y* observations in *E. coli* can be explained by a two-dimensional rate-yield tradeoff, where the first dimension is the characteristic *µ*–*Y* tradeoff associated with acetate overflow metabolism and the second dimension is a tradeoff between glucose uptake rate (*q_glc_*) and *Y* that appears during ALE adaptation. We employ a multi-scale modeling approach to provide a mechanistic description of the two-dimensional rate-yield tradeoff. By deriving the relationship between the ME-model and the previously reported small-scale proteome allocation model (Basan et al., 2015), we are able to develop a workflow for fitting ME-model parameters to experimental data, and we achieve quantitative accuracy for simulations of *µ*–*Y*. This multi-scale modeling approach predicts a two-dimensional rate-yield tradeoff, and it suggests that the second dimension of the tradeoff can be explained by changes in P/O ratio and the flux balance between glycolysis and pentose phosphate pathway.

## 2 RESULTS

### 2.1 Adaptive laboratory evolution reveals a two-dimensional rate-yield tradeoff

To explore the metabolic constraints on *E. coli* growth, adaptive laboratory evolution (ALE) was used to adapt *E. coli* K-12 MG1655 to maximize growth at 37°C in a liquid culture with a minimal medium containing glucose LaCroix et al. (2015). Eight independent experiments were performed on an automated ALE platform to achieve 8.3 × 10^12^ to 18.3 × 10^12^ cumulative cell divisions Lee et al. (2011). Phenotypic characterization was performed on eight ALE endpoint strains, including quantitative measurements of *µ*, *q_glc_*, *q_ac_*, and other common metabolic byproducts of *E. coli* (Methods).

A diversity of metabolic phenotypes was observed in the ALE endpoint strains. Through ALE, *µ* increased from 0.7 h^-1^ for wild-type (red triangles in Fig. 1A–D) to 0.95–1.10 h^-1^ (red circles with error bars in Fig. 1A–D). Based on previous reports, we expected a linear relationship between *µ* and *q_ac_*. However, ALE endpoint strains achieved a wide ranging *q_ac_* from 3.9–11.4 mmol gDW^-1^ h^-1^ (where wild-type *q_ac_* was 3.9 mmol gDW^-1^ h^-1^. While we did not observe a correlation between *µ* and *q_ac_* in these strains (Fig. 1B), there was a clear correlation between *q_glc_* and *q_ac_* (Fig. 1D).

These correlations have been observed previously for *E. coli* strains (LaCroix et al., 2015; Peebo et al., 2014; Bekker et al., 2009), and moreover, a bacterial engineering approach has been reported to vary *q_ac_* by manipulating the substrate uptake system (Lara et al., 2008). In one of these studies, Bekker et al. (2009) showed that switching electron transport chain (ETC) enzyme selection (and thereby modifying the P/O ratio) can cause a *q_glc_* –*Y* tradeoff at a low *µ* of 0.15 h^-1^. ALE gained *q_ac_* and *Y* decoupled from *µ*, which is seemingly contradict to the reported correlated *mu*–*Y* and *mu*–*q_ac_* (Basan et al., 2015; Nanchen et al., 2006). The ME-model we used in this study is aiming to simulate the relationships between these *q_glc_* –*q_ac_* and *q_glc_* –*Y* tradeoffs, connecting to the mechanisms of *µ*–*Y* tradeoffs (including Lipson’s bell-shaped curve, Fig. 1A) by established models (Basan et al., 2015; Mori et al., 2016).

To enable our analysis, it is important to note that ALE endpoint strains rapidly acquire regulatory mutations, but they do not acquire new metabolic capabilities within the time frame of these experiments (Sandberg et al., 2014; LaCroix et al., 2015; Utrilla et al., 2016). The linear correlation between *q_ac_* and *µ* reported previously was identified for an isogenic strain (Nanchen et al., 2006; Basan et al., 2015). In contrast, our observations of a decoupling between *q_ac_* and *µ* appear when comparing adapted strains. However, because these adapted strains have only regulatory mutations, their phenotypes represent the limits of what *E. coli* cells can achieve while bounded by metabolic and proteomic constraints (but not by regulation). This type of adaptation and the associated phenotypic tradeoffs are useful for understanding cellular adaptation to ecological niches where regulatory adaptation can occur rapidly (Elena and Lenski, 2003).

### 2.2 ME-model data fitting with a multi-scale modeling approach

To explain these experimental observations, we sought a modeling approach that could quantitatively predict the *µ*–*Y* and *µ*–*q_ac_* relationships. Our modeling approach starts with fitting the linear-threshold (blue line in Fig.1B) *mu*– *q_ac_* relation (Basan et al., 2015) using the framework of ME-model (O’Brien et al., 2013; Lloyd et al., 2018). We first considered a previously reported coarse-grained model of proteome allocation (Basan et al., 2015) that describes *E. coli* overflow metabolism (Fig. 1E). The *Y* -maximizing approach done by Basan et al. (2015) indicates that high-*Y* growth strategies have a fitness benefit in spatially structured environments (like biofilms) that has been demonstrated through a *Y* -selection system Bachmann et al. (2013), and more efficient strategies also leave more resources for cells that are hedging against future stresses Utrilla et al. (2016). The evolutionary history of *E. coli* includes growth in structured environments and a wide range of stresses that could have placed a selection pressure on increasing *Y*. Therefore, we focused on fitting the observed chemostat Nanchen et al. (2006) and uptake titration Basan et al. (2015) data for the *Y* -maximized growth solution (green and blue curves in Fig. 2).

**FIGURE 2.**
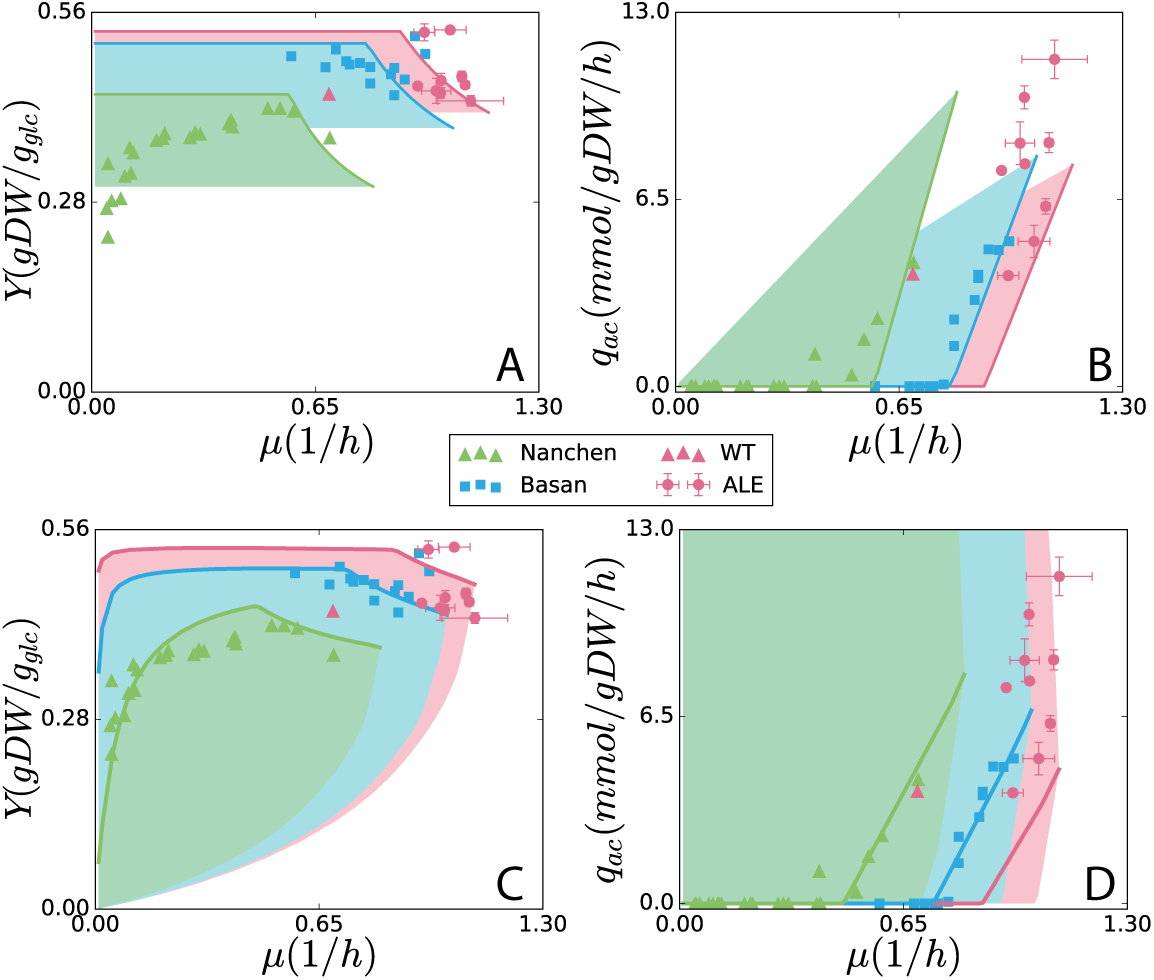
Growth phenotypes from simulations of *E. coli* (A, B) using the SSME-model and (C, D) using the ME-model. Simulations were fit to experimental data for each of the three datasets, for K-12 MG1655 chemostat (Nanchen et al., 2006) (green triangles)) and NCM3722 substrate titration (Basan et al., 2015) (blue squares)), and strains adapted from wild-type K-12 MG1655 (red triangle, LaCroix et al. (2015) for maximum growth rate through ALE (this study; red circles; error bars for standard deviation across duplicates). The *Y* -maximized solutions are displayed as solid lines in all plots. For both models, fitting was performed by manipulating three global parameters: unmodeled protein fraction (UPF), growth-associated maintenance (GAM), and non-growth associated maintenance (NGAM). Details of the fitting approach are provided in the Methods.

The coarse-grained proteome allocation model was intended to make predictions at high *µ* and thus only captures the negative *µ*–*Y* relation (Fig. 2A). The parameters in the coarse-grained model have a strong experimental basis in fine-grained protein abundances measurements in high growths, and the model produces accurate predictions of *µ*–*q_ac_* (Basan et al., 2015).

We also considered the genome-scale ME-model *i*JL678-ME (Lloyd et al., 2018). With the default parameter settings in the ME-model, simulations had a poor quantitative prediction (O’Brien et al., 2013) of *µ*–*q_ac_* to the uptake titration data (Fig. S3F). This poor fit can be explained by inaccurate genome-wide enzyme turnover rates (*k_eff_* s) that ME-model researchers have been seeking to improve (Lloyd et al., 2018; Ebrahim et al., 2016; Nilsson et al., 2017). We sought to modify the *k_eff_* s to fit the *µ*–*q_ac_* data. However, since each of the 5266 reactions in the genome-scale ME-model has a *k_eff_* parameter, it is difficult to directly fit the parameters to measured data.

Therefore, we pursued a multi-scale modeling approach where the coarse-grained model was used to analyze the effects of proteome-efficiency at the level of complete pathways, and this insight was used to tune parameters in the ME-model in bulk. To connect the models, we first found that the proteome efficiency (*ε*) parameters in the coarse-grained model share a conceptual basis with the enzyme efficiency parameter *k_eff_* s in ME-models (“2 Proteome constraints in the ME-model” in Appendix). Thus, we were able to reformulate the coarse-grained model within the framework of a ME-model (Fig. S1). The resulting small-scale ME-model (SSME-model) has parameters directly analogous to those in the genome-scale ME-model (See “4 SSME-model parameters derivation” and “5 Matlab and COBRAme implementation” in Appendix). The resulting SSME-model generates identical *µ*–*Y* and *µ*–*q_ac_* predictions to the proteome allocation model. The most obvious difference between the SSME-model derived from Basan et al. (2015) model and ME-model for these phenotypic predictions is the expanded solution space of the ME-model (Fig. 2). However, much of the ME-model solution space corresponds to very low yield metabolic solutions. If *Y* is maximized during simulations of the SSME-model and ME-model (achieved by minimizing *q_glc_* at a given *µ*), the resulting predictions are more similar between the models and lie closer to experimental data (solid blue curves in Fig. 2).

To enable ME-model fitting starting from the poor fit in Fig. S3F, we first analyzed parameter sensitivities in the SSME-model. The SSME-model has a small number of *k_eff_* s, with only three pathways (respiration, fermentation, biomass) with *k_eff_* s, making it easy to test the sensitivities of predictions to changes in *k_eff_* (Cheng, 2017). The SSME-model shows that dropping the *k_eff_* for respiration decreases the slope of the *µ*–*q_ac_* line without affecting *µ*_*max*_, suggesting that dropping the *k_eff_* s of TCA cycle enzymes in genome-scale ME-model would lead to a more accurate fit (“7 Experimental data fitting” in Appendix). However, only dropping the TCA *k_eff_* s leads to a *µ*–*q_ac_* slope that is still not gradual enough (Cheng, 2017). Next, we determined that secondary pathways in the ME-model carry unrealistic fluxes, and they could be responsible for the remaining prediction gap. We used an iterative approach to block the metabolic reactions in the ME-model (“7 Experimental data fitting” in Appendix) and achieved an accurate ME-model prediction of the *µ*–*q_ac_* acetate line (solid blue line in Fig. 2D and Fig. S3F). The three global parameters unmodeled protein fraction (UPF), growth-associated maintenance (GAM), and NGAM (Table S2 in Appendix) were then used to fit predictions to individual datasets (green, blue and red curves in Figure 2).

### 2.3 The ME-model predicts phenotypic diversity in ALE strains

As a result of data fitting, we achieved a quantitative fit of chemostat (Nanchen et al., 2006) and uptake titration (Basan et al., 2015) data with the *Y* -maximized ME-model solutions (blue and green curves in Fig. 2C,D). The ALE-adapted strains (red circles in Fig. 2) do not align well with the *Y* -maximizing solutions (red curves in Fig. 2), but they are encompassed by the ME-model solution space, and further analysis of these data points and the corresponding ME-model solutions were used to understand the phenotypic diversity of these adapted strains.

Feasible solutions other than the*Y* -maximized solution are achieved through the activation of alternative metabolic pathways. The SSME-model does not capture the ALE data points with high *q_ac_* (red region in Fig. 2B), while the genome-scale ME-model does (red region in Fig. 2D). Moreover, the ME-model predicts feasible growth at lower *Y* in the *µ*–*Y* solution space than the SSME-model. We sought to determine which pathways are responsible for the lower *Y* and higher *q_ac_* in ME-model that was not captured by the SSME-model.

Removing reactions from the ME-model can decrease the size of the solution space (Fig. S4, “8 Solution space variation” in Appendix), making the solution space more similar to the SSME-model solution space. We employed a workflow to identify 24 reactions (Table S3 in Data) that are not activated in the *Y* -maximized solutions but are used to enable higher *q_ac_* at lower *Y*. We observed that these 24 reactions are part of metabolically inefficient pathways that are alternatives to the high *Y* pathways in *Y* -maximized solutions. By extension, metabolically inefficient pathways can be added to the SSME-model to increase the size of the solution space (Fig. 10), making it more similar to the ME-model solution space. Thus, the modified SSME-model can achieve low *Y* (Fig. 10A) at high *q_ac_* (Fig. 10C). Therefore, the difference in predictions between the ME-model and SSME-model is a result of the greater range of metabolic capabilities of the genome-scale model.

### 2.4 The two-dimensional rate-yield tradeoff

We can now provide a theory to connect the correlations in *µ*–*Y* (Fig. 1A) (and the associated acetate curve in *q_ac_* –*Y*, Fig. 1B) with the negative correlation in *q_glc_* –*Y* (Fig. 1C) and positive *q_glc_* –*q_ac_* correlations (Fig. 1D).

To see the relationship between the three variables *µ, q_glc_*, and *Y* we generated ME-model solution spaces in *q_glc_* and *Y* at increasing lower bounds of *µ* (Fig. 3A). These solution spaces represent the flexibility in the model to achieve a growth rate. At the *Y* -maximed limit of these solution space, we see the established negative *µ*-*Y* tradeoff where increasing growth rate requires decreasing *Y* (dashed arrow marked as “d1” in Fig. 3A) coupling with increasing *q_ac_* (top edges of solution spaces in Fig. 3B). This is the first dimension of the rate-yield tradeoff.

**FIGURE 3.**
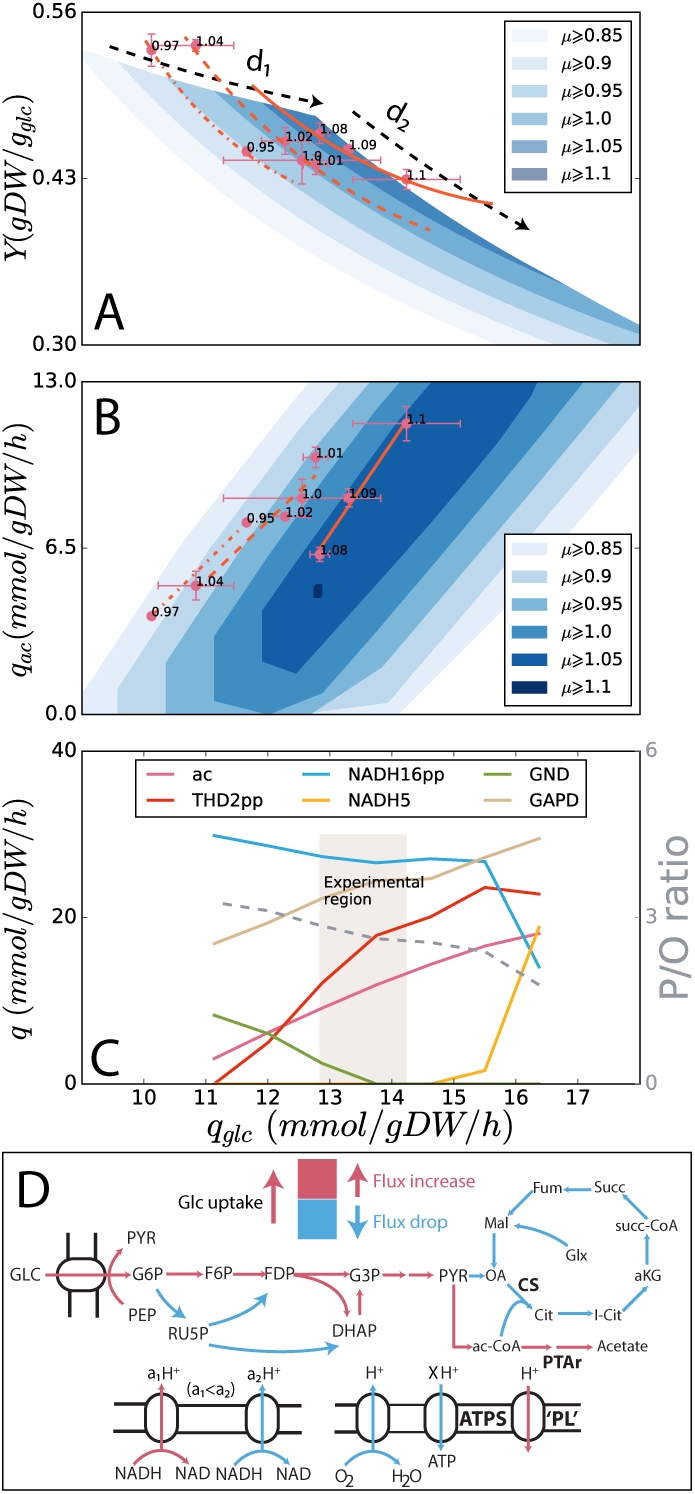
Analysis of the second dimension of the rate-yield tradeoff. (A) Two dimensions of the rate-yield tradeoff. The first dimension “d1” is the negative *µ*–*Y* correlation at maximum *Y*, and the second dimension “d2” is the negative *q_glc_* –*Y* correlation at a fixed *µ*. These correlations are observed in ME-model simulations and experimental data from ALE strains. (B) A correlation between *q_glc_* and *q_ac_* is also observed at fixed *µ* in both the ME-model and ALE endpoint data. Linear fits for the experimental data at similar growth rates are shown as dash-dotted (*µ*=0.95–0.97 h^-1^, dashed (*µ*=1.00–1.04 h^-1^, and solid (*µ*=1.08–1.10 h^-1^ orange lines. These fits are described by the upper edges of the *q_glc_* –*q_ac_* solution space at fixed *µ*. For growth between 1.00 and 1.04 h^-1^, *r* ^2^=0.931 and *p*=0.035. For growth between 1.08 and 1.1 h^-1^, *r* ^2^=0.986 and *p*=0.071. (C) The reaction fluxes in ME-model simulations along the upper edge (maximizing *q_ac_*) of the solution space for *µ*=1.05 h^-1^. Notably, the P/O ratio (gray dashed curve) is decreasing with increasing *q_glc_*. (D) A pathway map of central metabolism showing the changes of the reaction fluxes with increasing *q_glc_* and *q_ac_*. “a1” and “a2” represent the rate of proton flux through the membrane. **Abbreviations. ac**: acetate excretion; **THD2pp**: NAD(P) transhydrogenase (catalyzed by the gene product of *pntAB*; **NADH16pp**: NADH dehydrogenase (*nuoA–N*; **NADH5**: NADH dehydrogenase (*ndh*; **GND**: Phosphogluconate dehydrogenase (*gnd*; **GAPD**: Glyceraldehyde-3-phosphate dehydrogenase (*gapA*; **CS**: Citrate synthase (*gltA*; **PTAr**: Phosphotransacetylase (*pta* and *eutD*; **ATPS**: ATP synthase (*atpA–I*; **PL**: model reaction representing proton leakage (Methods).

At a given *µ*, the ME-model solution spaces extend toward lower*Y* and higher *q_glc_*, revealing an inverse proportional relationship in *q_glc_* -*Y*. This relationship is also observed in ALE endpoint strains with similar *µ* (Fig. 3A). This is the second dimension of the rate-yield relationship (“d2” in Fig. 3A) defining the second-order rate-yield tradeoff. The second dimension can also be seen in *q_ac_* –*q_glc_* where the ME-model predicts the *q_ac_* –*q_glc_* correlation observed in ALE endpoint as the *q_ac_* -minimized edge of the solution space (Fig. 3B). Interestingly, the solution spaces predicted by ME-model show broad feasible ranges of acetate production at a given *q_glc_* and *µ* (“bold” solution spaces in Fig. 3B), so the *q_glc_* –*q_ac_* tradeoff is not required by the model. On the other hand, the relationship between *q_glc_* and *Y* is a strict tradeoff in the model (“thin” solution spaces in Fig. 3A), so we propose that *q_glc_* –*Y* is the more fundamental second dimension of the rate-yield tradeoff.

### 2.5 Mechanisms for the additional rate-yield tradeoff

We sought to identify the particular alternate metabolic strategies in the ME-model that could enable a *q_glc_* –*Y* tradeoff by identifying the differential pathway usage at a fixed high *µ* (1.05 h^-1^ in the ME-model (Fig. 3C). The model predicts that when *q_glc_* increases from the *Y* -maximized state (minimum *q_glc_*), flux through the proton-coupled NAD(P) transhydrogenase increases (reaction THD2pp, catalyzed by *pntAB*. In addition, a pathway switch between two different NADH dehydrogenase reactions, NADH5 (*ndh* and NADH16pp (*nuo*, appears at high *q_glc_*. In fact, each of or any combination of the 24 reactions in Table S4 (Expanded view, Data) can be activated in the ME-model to achieve high *q_glc_*, high *q_ac_*, and low *Y*. There are two common threads among these pathway activations. First, they all decrease the P/O ratio in the simulations (Fig. 3C). NADH5 contributes fewer protons to the periplasm per electron than NADH16pp. And increasing THD2pp flux drains the proton gradient without contributing to ATP production, thereby decreasing P/O ratio (Fig. 3D). Second, glycolytic flux increases (Fig. 3D) and pentose phosphate pathway flux decreases (Fig. 3C).

Experiments that introduce proton leakage have shown a shift towards high *q_ac_* and low *Y* (Basan et al., 2015). It has also been shown that the variation of P/O ratio can uncouple the regulation of cytochrome oxidase from the cellular ATP demand (Bekker et al., 2009). More broadly, energy dissipation through proton leakage is known to be a method of metabolic control in bacteria (Russell and Cook, 1995; Russell, 2007). To clarify the effect of decreasing of P/O ratio in the ME-model, we added a reaction in the model representing proton leakage (Methods). As a result, we see the *Y* -maximized solution with decreased P/O ratios have higher *q_glc_*, higher *q_ac_*, and lower *Y* at a given *µ* (Fig. 4). Finally, experiments have shown that knocking out *gnd* leads to increased *q*_*g*__*l* *c*_ and *q_ac_* and decreased *Y* with little change in *µ* (Jiao et al., 2003). The ME-model also predicts that *gnd* knockout mutants (“gnd knockout simulation” Methods) will have increased *q_glc_*, *q_ac_* and decreased *Y* (Fig. 4). Thus, the ME-model points to general mechanisms for this fundamental second-order tradeoff, but the exact pathways involved can be determined in future experiments, and it may be that multiple pathways work together to enable it.

**FIGURE 4.**
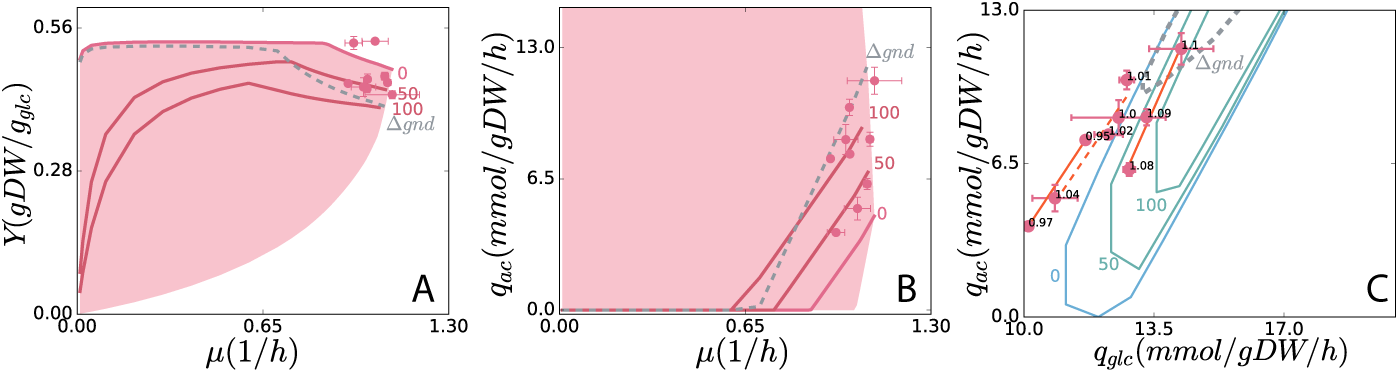
The second-order rate-yield tradeoff demonstrated by decreasing the P/O ratio and and knocking out *gnd* (“Δ*gnd*”) in ME-model simulations. The drop of P/O ratio is achieved by inducing the proton leakage (“PL” in Fig. 3D) reaction flux, as 0, 50 mmol gDW^-1^ h^-1^ (labeled “50”), and 100 mmol gDW^-1^ h^-1^ (labeled “100”). The new *Y* -maximized solution curves (solid red for “PL” flux variation, dashed grey for Δ*gnd*) and the *q_ac_* –*q_glc_* solution space contours (fixed *µ*=1.0 h^-1^, solid blue for “PL” flux variation, dashed grey for Δ*gnd*) were simulated in the ME-model.

Alternative explanations of the rate-yield tradeoff have been proposed, including membrane (Zhuang et al., 2011; Szenk et al., 2017) and cytosolic crowding (Adadi et al., 2012; Vazquez and Oltvai, 2016). It is difficult to rule out these alternative constraints on cell growth, and it may be that multiple constraints operate at once. However, it is encouraging to see that the ME-model can explain the complex relationship between *µ, Y, q*_*ac*_, and *q_glc_* with only metabolic and proteome allocation constraints. In the future, it will be possible to extend ME-models with additional constraints. For example, it has been proposed that the UPF parameter is growth-rate dependent, and thus existing proteome allocation models with fixed UPF are inaccurate (Vazquez and Oltvai, 2016). If this is indeed the case, then SSME- and ME-models with cytosolic crowding constraints can be developed to fully represent the interplay between crowding, proteome allocation, and pathway selection.

## 3 DISCUSSION

The *E. coli* ME-model provides a mechanistic and predictive model of rate-yield tradeoffs. It successfully reconciles several experimental data sets: i) uptake titration at low growth (Nanchen et al., 2006), ii) batch culture at higher growth rates (Basan et al., 2015), and iii) ALE endpoint strains (this study). These data sets, when analyzed with the ME-model, show the existence of a two-dimensional rate-yield tradeoff. This two-dimensional tradeoff cannot be deciphered from simpler intuitive models, but it can be derived from the comprehensive set of biochemical mechanisms represented by the ME-model.

Furthermore, this study employed a multi-scale modeling approach where a small-scale model was used to guide parameter estimation in the genome-scale ME-model. This approach—which has been termed Tunable Resolution (TR) modeling (Kirschner et al., 2014)—was essential to the success of the study, and we expect that both small-scale and genome-scale models will continue to play an important role in understanding the genotype-phenotype relationship.

The two-dimensional rate-yield tradeoff appears as a result of ALE selection for *µ* when alternative pathway selection strategies achieve the same growth rate. Proton leakage and alternative ETC pathway selection are plausible mechanisms for modifying the P/O ratio and creating the *q_glc_* –*Y* tradeoff. In addition, the flux ratio between glycolysis and the pentose phosphate pathway might play a significant role in the *q_glc_* –*Y* tradeoff. Those mechanisms can be tested experimentally. Finally, revealing the underlying regulation would be of great interest for establishing a deeper understanding of rate-yield tradeoffs. Combining ME-models with known regulatory mechanisms to explain cellular choices would achieve a long-standing goal in systems biology (Reed and Palsson, 2003).

## 4 MATERIALS AND METHODS

Phenotypic data including *µ, q_glc_*, *q_ac_*, and excretion rates of other metabolic byproducts were collected for ALE endpoint strains (“1 Phenotypic characterization of *E. coli* strains” in Appendix). Reference data points were collected from published studies Nanchen et al. (2006); Basan et al. (2015). The coarse-grained proteome allocation model from Basan et al. (2015) was reformulated as a small-scale ME-model (SSME-model, detail in “4 SSME-model parameters derivation” in Appendix) and implemented by the COBRAme framework Lloyd et al. (2018). The genome-scale model *i*JL1678-ME was modified to fit experimental data by modifying the *k_eff_* s of TCA cycle reactions, blocking target reactions, and modifying UPF, GAM, and NGAM (“7 Experimental data fitting” in Appendix). Solution spaces were generated using flux balance analysis (incorporated in COBRAme) in the ME-model (“6 Solution space of the ME-model” in Appendix). To determine the effect of modifying P/O ratio on ME-model solution spaces, a reaction representing proton leakage was added to the ME-model (“9 P/O ratio manipulation” in Appendix). The effect of the *gnd* knockout was demonstrated by blocking the reaction GND in ME-model simulations (“10 gnd knockout simulation” in Appendix).

## EXPANDED VIEW

### Appendix

The appendix includes the detailed introduction and discussions of the materials and methods.

### Data

Includes Tables S3–S10. Table S3 presents the blocked reactions in the ME-model to achieve a quantitative fit to experimental data. Table S4 shows the essential exchange reactions that should be turned on for maintaining growth. Tables S5 and S6 show the target reactions that vary the solution space of the ME-model. Tables S7–S10 include all data that are presented in the figures in the paper.

## ACKNOWLEDGEMENTS

The authors would like to thank Ali Ebrahim, Laurence Yang, and Colton Lloyd for the assistance with ME-model analysis. Moreover, we are grateful to Ke Chen, David Heckmann, Marc Abrahms, Amitesh Anand and Brian Taylor for providing their advice on the manuscript. We highly appreciate Andrea De Martino for providing single strain (K-12 MG1655 and NCM3722) growth yield and acetate overflow data. And at last, we would like to express our extraordinary thanks to Terence Hwa and Matteo Mori for advice on developing multi-scale models of proteome allocation.

## AUTHORSCONTRIBUTIONS

A.M.F, B.O.P, and Z.A.K designed research; C.C, E.J.O, and Z.A.K performed research; D.M, J.U, C.O, R.A.L, and T.E.S generated and analyzed experimental data; C.C, C.O, B.O.P, and Z.A.K wrote the paper.

## CONFLICTOFINTEREST

The authors declare no conflict of interest.

